# Optimal AAV capsid/promoter combinations to target specific cell types in the common marmoset cerebral cortex

**DOI:** 10.1101/2024.05.09.593444

**Authors:** Yasunori Matsuzaki, Yuuki Fukai, Ayumu Konno, Hirokazu Hirai

## Abstract

To achieve cell type-specific gene expression, using target cell-tropic AAV capsids is advantageous. However, their tropism across brain cell types remains unexplored in non-human primates. We assessed the tropism of nine AAV serotype capsids (AAV1, 2, 5, 6, 7, 8, 9, rh.10 (rh10), and DJ) on marmoset cerebral cortical cell types. Marmoset cerebral cortex was injected with different serotype AAVs expressing enhanced GFP (EGFP) by the ubiquitous chicken β-actin hybrid (CBh) promoter. After 4 weeks, all nine AAV capsid vectors, especially AAV9 and AAVrh10, caused highly neuron-selective EGFP expression. Some AAV capsids, including AAV5, caused EGFP expression in oligodendrocytes to a lesser extent, with minimal or no expression in astrocytes and microglia. Different ubiquitous CMV and CAG promoters showed similar neuron-predominant transduction. Conversely, all nine AAV capsid vectors with the astrocyte-specific hGFA(ABC1D) promoter selectively transduced astrocytes, except AAV5, which transduced oligodendrocytes modestly. Oligodendrocyte-specific mouse myeline basic protein (mMBP) promoter in AAV5 vectors transduced oligodendrocytes specifically and efficiently. Our results suggest optimal combinations of capsids and promoters for cell type-specific expression: AAV9 or AAVrh10 and ubiquitous CBh, CMV, or CAG promoter for neuron-specific transduction; AAV2 or 7 and hGFA(ABC1D) promoter for astrocyte-specific transduction; and AAV5 and mMBP promoter for oligodendrocyte-specific transduction.

## Introduction

The brain is an organ in which an extremely large number of cells, including neurons, astrocytes, oligodendrocytes, and microglia, extend their processes and are intricately intertwined.

Recent studies have shown that specific cell types play important roles in the onset and progression of brain diseases, including oligodendrocytes in multiple sclerosis ^1,2^, microglia in Alzheimer’s disease ^3,4^, and astrocytes and microglia in stroke ^5,6^. For these brain diseases, delivering and expressing genes to specific cell types involved in pathogenesis may allow to elucidate the underlying molecular mechanisms and develop therapeutic interventions. However, unintended expression of a gene in non-target cell types can cause non-specific effects, which makes result interpretation difficult and, in case of gene therapy, potentially leading to adverse events.

To express a gene specifically and efficiently in a target cell type, using a target cell-tropic capsid and a target cell-selective promoter is crucial. Although the tropism of different serotype capsids has been studied in rodents ^7,8^, it has not been well studied in non-human primates, probably due to the limited number of animals available.

In this experiment, we aimed to explore the tropism of nine different serotype capsids to distinct brain cell types in the non-human primate, common marmoset and to examine whether combination of target cell-tropic capsids with appropriate promoters enables target cell-specific gene expression in marmoset brain.

## Results

To compare the cellular tropism of different AAV serotypes in the marmoset brain, we injected nine AAV serotypes (AAV1, 2, 5, 6, 7, 8, 9, rh.10 (rh10), and DJ) into the marmoset brain (see Table 1 for marmoset details). The AAV vectors were designed to express EGFP under the control of the constitutively active chicken β-actin hybrid (CBh) promoter (Figure 1A). Each AAV vector was injected into up to 10 locations in the marmoset cerebral cortex (Figure 1B). Since injection of high titer AAV causes inflammation and local tissue damage ^9^, we first determined the optimal viral titer for the assessment. The marmoset cerebral cortex was injected with a triple dilution series of the AAV2 vector (1 μL/each point). EGFP expression and tissue conditions were examined 4 weeks after the viral injection by fluorescent immunohistochemistry (fIHC). Low magnified EGFP fluorescence images showed that the EGFP labeling area and fluorescence intensity were roughly proportional to the injected viral titer without affecting NeuN labeling (Figure S1A), showing no neuronal loss over the range of AAV titers used. However, in the area that received the highest viral titer (2.0 × 10^9^ vg), microglia shape changed markedly with increased immunoreactivity, and microglia processes surrounded the neuronal cell bodies (Figure S1B). These results suggest a strong inflammatory response and tissue damage from the injection of high doses of AAV.

**Figure 1.**
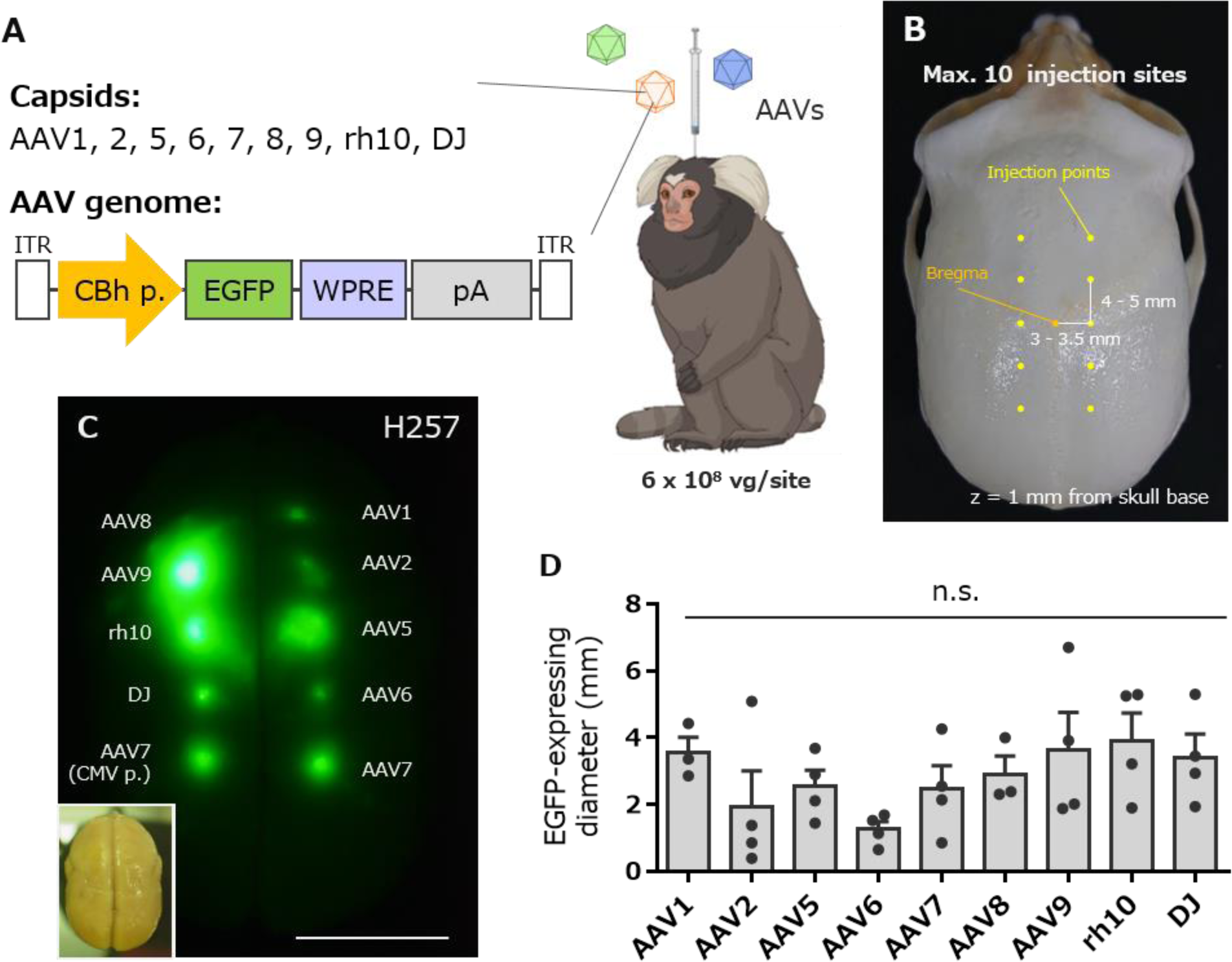
Comparison of expression levels of EGFP by injection of nine different serotype AAV vectors into the marmoset cortex. (**A**) Schema depicting injection of AAV vectors to a marmoset. Nine AAV capsid vectors expressing EGFP under the control of the CBh promoter were injected into the marmoset cerebral cortex. (**B**) The coordinate of the viral injection with reference to the bregma of the marmoset skull. (C) Example of EGFP fluorescence image of marmoset cortex 4 weeks after viral injection. The lower left inset is a bright field brain image of a marmoset (Animal ID: H257, see Table 1). Scale bar, 10 mm. (D) Graph showing the diameter of EGFP fluorescence on the cortex. No statistically significant differences were observed in the EGFP fluorescence diameters by one-way ANOVA with Tukey’s post hoc test. n.s., not significant.

**Table 1.** Marmoset profiles used for analysis of nine serotypes of AAV vectors expressing EGFP by CBh, CMV, CAG, hGFA(ABC1D) and mMBP promoter.

| ID | Name | Sex | AAV vector injection |  |  |  |  | Sacrifice |  |  |
| --- | --- | --- | --- | --- | --- | --- | --- | --- | --- | --- |
| | | | Promoter | Titer<br>(vg/<br>serotype) | Volume<br>(μL) | Old<br>(y) | BW<br>(g) | Incubation<br>time<br>(days) | BW<br>(g) | Weight<br>change<br>(fold) |
| H227 | Shinobu | Male | CBh | $6.0 \times 10^8$ | 1 | 1.6 | 335 | 32 | 316 | 0.94 |
| H231 | Hamo | Male | CBh | $6.0 \times 10^8$ | 1 | 1.4 | 365 | 34 | 374 | 1.02 |
| H232 | Odoru | Male | CBh | $6.0 \times 10^8$ | 1 | 1.3 | 325 | 34 | 339 | 1.04 |
| H257 | Nishin | Male | CBh, CMV | $6.0 \times 10^8$ | 1 | 1.1 | 320 | 28 | 314 | 0.98 |
| H259 | Shirasu | Female | CBh, CMV | $6.0 \times 10^8$ | 1 | 1.1 | 338 | 29 | 340 | 1.01 |
| H260 | Hirame | Male | CBh, CMV | $6.0 \times 10^8$ | 1 | 1.1 | 384 | 29 | 391 | 1.02 |
| H263 | Maru | Male | CBh, CMV | $6.0 \times 10^8$ | 1 | 1.1 | 408 | 28 | 399 | 0.98 |
| H149 | Sango | Female | CAG | $6.0 \times 10^8$ | 1 | 5.8 | 340 | 28 | 357 | 1.05 |
| H276 | Yo | Female | CAG | $6.0 \times 10^8$ | 1 | 2.6 | 505 | 31 | 494 | 0.98 |
| H309 | Azami | Female | CAG | $6.0 \times 10^8$ | 1 | 1.7 | 488 | 29 | 487 | 1.00 |
| H312 | Hijiki | Male | CAG | $6.0 \times 10^8$ | 1 | 1.7 | 529 | 28 | 527 | 1.00 |
| H256 | Sawara | Male | hGFA(ABC1D) | $6.0 \times 10^8$ | 1 | 1.4 | 371 | 31 | 395 | 1.06 |
| H268 | Noko | Female | hGFA(ABC1D) | $6.0 \times 10^8$ | 1 | 1.2 | 316 | 29 | 328 | 1.04 |
| H269 | Take | Male | hGFA(ABC1D) | $6.0 \times 10^8$ | 1 | 1.2 | 344 | 33 | 401 | 1.17 |
| H271 | Aoba | Male | hGFA(ABC1D) | $6.0 \times 10^8$ | 1 | 1.2 | 333 | 29 | 344 | 1.03 |
| H112 | Azuki | Female | mMBP | $6.0 \times 10^8$ | 1 | 7.0 | 354 | 33 | 372 | 1.05 |
| H262 | Madoka | Female | mMBP | $6.0 \times 10^8$ | 1 | 2.3 | 450 | 30 | 467 | 1.04 |
| H291 | Kanro | Male | mMBP | $6.0 \times 10^8$ | 1 | 1.7 | 511 | 32 | 444 | 0.87 |
| H295 | Santa | Male | mMBP | $6.0 \times 10^8$ | 1 | 1.5 | 362 | 28 | 355 | 0.98 |

Since such pathological changes were not observed at AAV doses less than 6.0 × 10^8^ vg (Figure S1C), we decided to use 6.0 × 10^8^ vg of AAVs for the following experiments. Nine different serotype vectors were injected into the cerebral cortex of marmosets, sacrificed 4 weeks later, and the size of the GFP-expressing area and transduced cell types were analyzed. Diameters of GFP fluorescent area were measured from the cerebral surface to compare the strength of expression of each capsid (Figure 1C). The diameters of GFP fluorescence were not significantly different among the nine AAV serotypes (Figure 1D; n = 3–4 marmosets, *p* = 0.26 by one-way ANOVA with Tukey’s post hoc test); however, a tendency indicating that the GFP fluorescent areas upon AAV2 and AAV6 injection were smaller can be observed (Figure 1C, D).

Next, we prepared cerebral cortical sections to examine cell types expressing GFP. The proportion of EGFP-expressing cell types (neurons, astrocytes, oligodendrocytes, and microglia) to total transduced cells was examined by fIHC. Because it was difficult to simultaneously immunolabel four different cell marker proteins in a single cerebral section, two serial sections were used: one immunostained with antibodies against NeuN, a neuron marker, and glial fibrillary acidic protein (GFAP) or S100β ^10^, astrocyte markers, and the other with antibodies against Olig2, an oligodendrocyte marker, and Iba1, a microglial marker (Figure 2A). An astrocyte marker, GFAP, is a membrane protein expressed primarily in the astrocyte processes, and its expression levels increase depending on tissue damage and inflammation ^11,12^. To measure the density of astrocytes in the intact cortex, we used S100β ^10^ instead of GFAP, as faint GFAP immunolabeling makes it difficult to identify astrocyte cell bodies.

**Figure 2.**
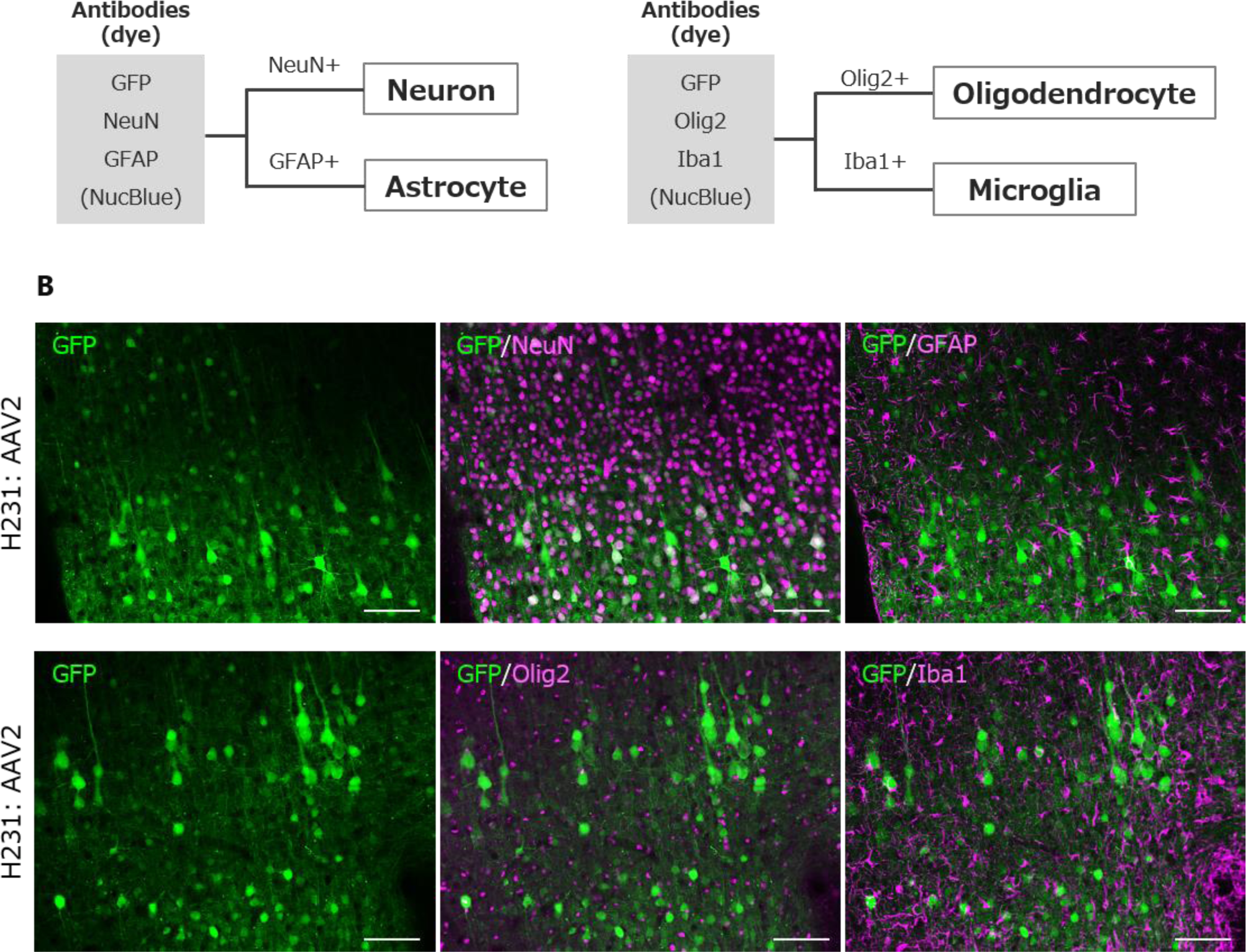
Identification of cell types by fluorescent immunohistochemistry (fIHC) (**A**) Scheme to identify EGFP-expressing cell types by fIHC. Neurons and astrocytes were immunolabeled for NeuN and GFAP, respectively. Cells immunostained for Olig2 or Iba1 were identified as oligodendrocytes and microglia. Cells were detected with Hoechest 33342, NucBlue, in the mounting reagent ProLong Glass. (**B**) Representative immunohistochemical images of marmoset cerebral cortex that received injection of AAV2. Two serial slices were presented: one immunolabeled for GFP, NeuN, and GFAP, and another one for GFP, Olig2, and Iba1, as indicated at each panel. Scale bar, 100 μm.

First, we measured the percentage of cell types present in the intact cerebral cortex of marmosets by immunohistochemistry. The cerebral cortex close to the injected sites (parietal lobe, Brodmann’s area 7) contained 43.4% of neurons, 11.7% of astrocytes, 41.9% of oligodendrocytes, and 3.0% of microglia (counted area 0.67 mm^2^, n = 3 marmosets analyzed, Figure S2). If AAV vectors unbiasedly infect and transduce cortical cells, the proportion of cell types expressing EGFP should follow the proportion endogenously present. However, the results showed that approximately 80–95% of EGFP-expressing cells were neurons in all AAV serotypes examined (Figure 3A, B). Notably, nearly all EGFP-expressing cells were neurons when injected with AAV2, 6, 9, rh10, or DJ, whereas AAV5 showed a significantly lower ratio of neurons to total EGFP-expressing cells compared with the highly neurotropic serotypes (Figure 3B; n = 3–4 marmosets, \*\**p* <0.01, \**p* <0.05 by one-way ANOVA with Tukey’s post hoc test).

**Figure 3.**
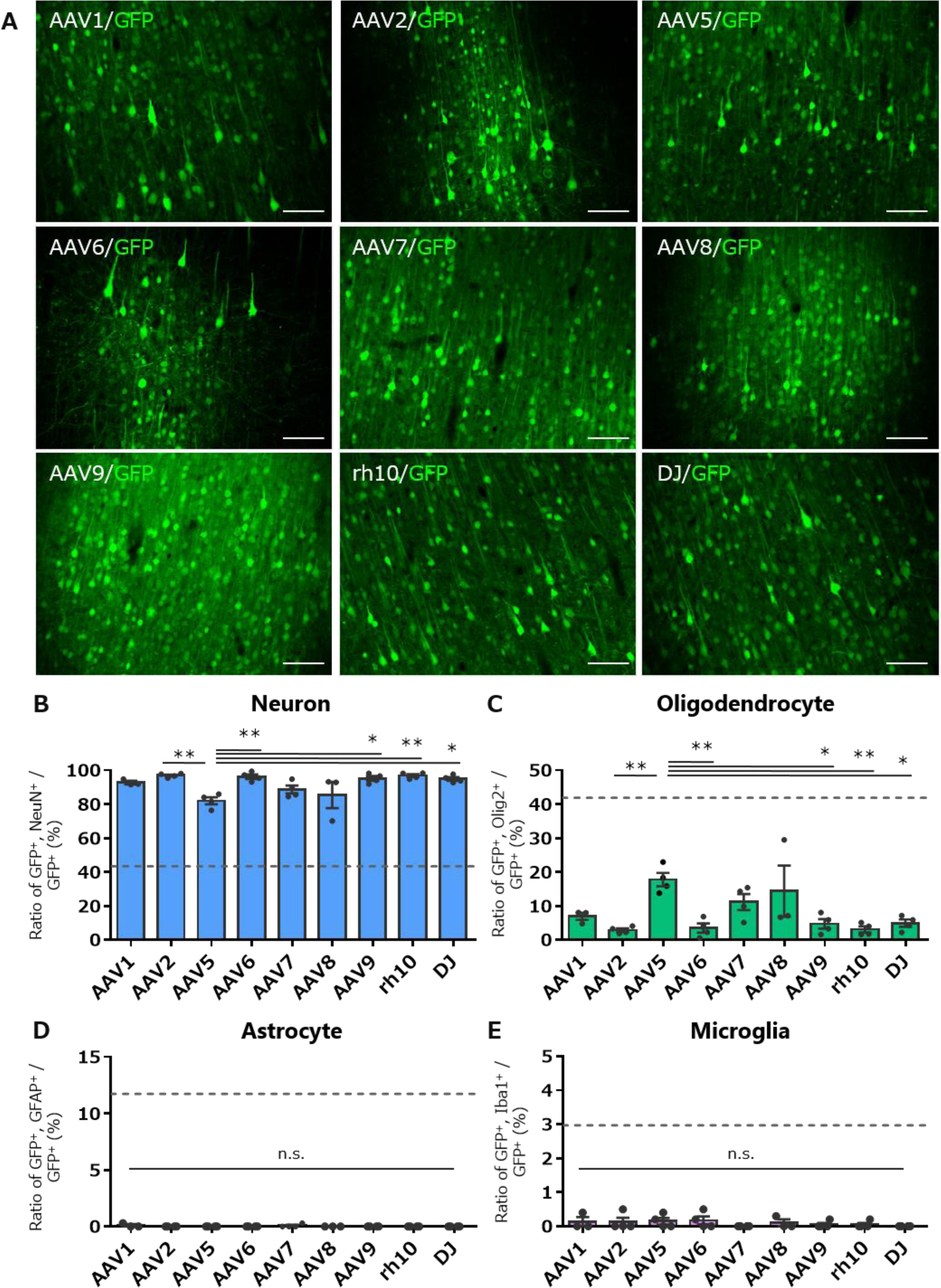
Quantitative analysis of transduced cell types following injection of nine serotype AAV vectors into the marmoset cerebral cortex. (**A**) Fluorescent images of the cortex immunolabeled for EGFP 4 weeks after AAV injection. Scale bar, 100 μm. (B-E) Graphs showing the ratio of GFP (+) neurons (**B**), GFP (+) oligodendrocytes (**C**), GFP (+) astrocytes (**D**), and GFP (+) microglia (E) to total GFP (+) cells. The dotted lines in the graphs are ratios of respective cell types to total cells present in the marmoset cortex (Figure S3). Error bars indicate S.E.M., and the dots in the graph indicate the respective values for each of the individual marmosets. The asterisks indicate a statistically significant difference between the AAV5 vector and the other capsids. \**p* < 0.05, \*\**p* < 0.01 by 1-way ANOVA with Tukey’s post hoc test. n.s., not statistically significant.

A minor fraction of EGFP-expressing cells were olig2-labeled oligodendrocytes, in which AAV5 transduced a significantly higher proportion of oligodendrocytes than many other serotypes (Figure 3C; n = 3–4 marmosets, \*\**p* <0.01, \**p* <0.05 by one-way ANOVA with Tukey’s post hoc test). However, the ratio of oligodendrocytes to all EGFP-expressing cells (∼20%) was much lower than the proportion of oligodendrocytes to the cells present in the cerebral cortex (42%). In contrast, none of the AAV serotypes tested transduced astrocytes or microglia (Figure 3D, E). Thus, all 9 AAV serotype vectors tested preferentially transduced neurons in the marmoset cerebral cortex, and AAV5 has the characteristic of transducing oligodendrocytes more than many other serotypes.

The lack of EGFP expression in astrocytes and microglia could be explained by a loss of CBh promoter activity in these cell types. Therefore, we performed similar experiments using AAV7 expressing EGFP under the control of another constitutive promoter, the CMV or CAG promoter. However, essentially the same results as seen with the CBh promoter were obtained. Injection of AAV7 vectors carrying the CMV or CAG promoter primarily transduced neurons and modestly oligodendrocytes, and no or little astrocytes and microglia were transduced (Figure 4). The transduction ratios of respective cell types did not show significant differences among the three promoters (Figure 4B-E; n = 4 marmosets, one-way ANOVA with Tukey’s post hoc test). Although not statistically significant, the CAG promoter showed a tendency to transduce astrocytes more than the CBh and CMV promoters (Figure 4C).

**Figure 4.**
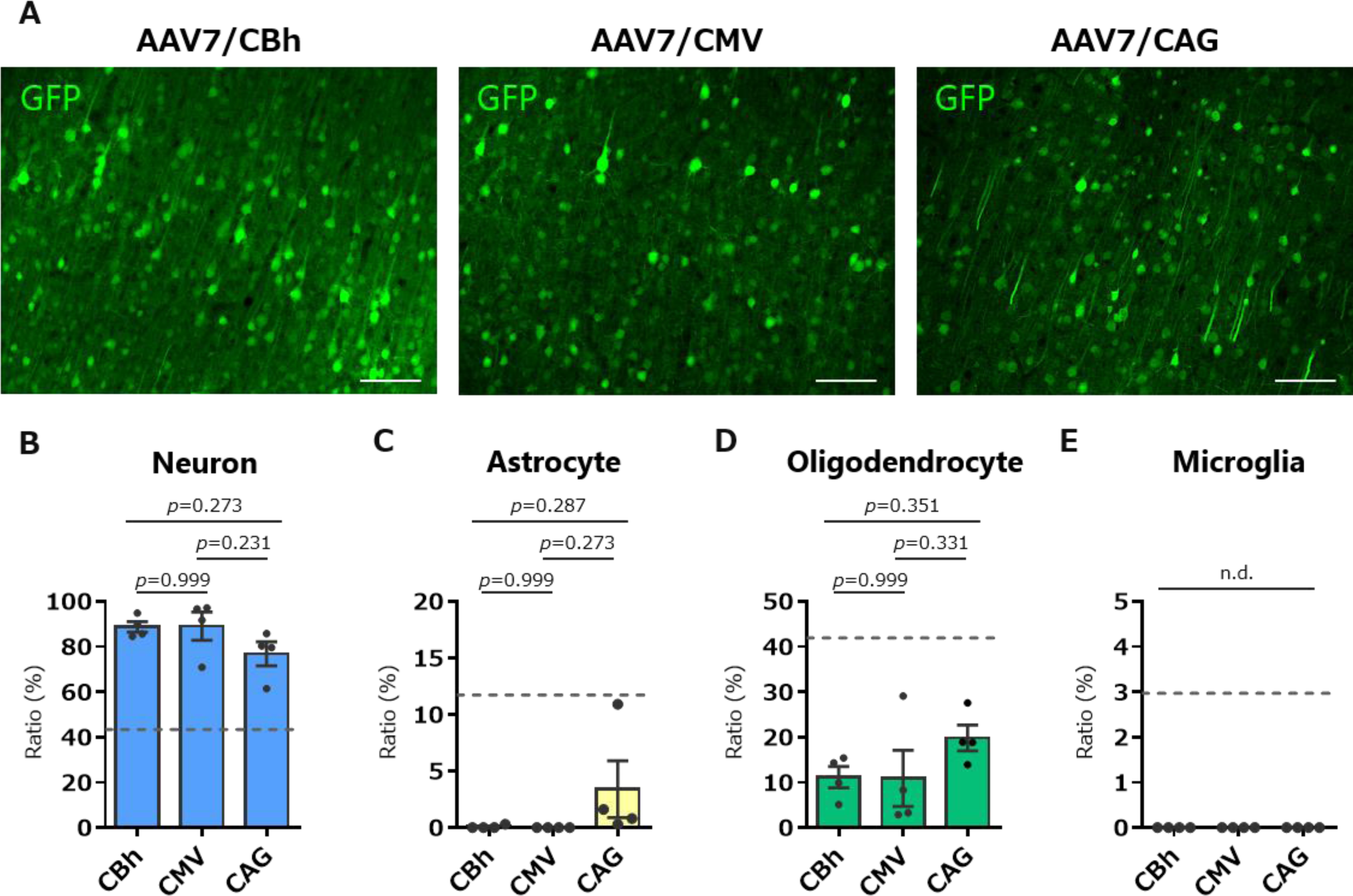
No or little glial cell transduction in the marmoset cortex by AAV vectors expressing EGFP under the control of 3 different ubiquitous promoters. (**A**) Immunofluorescent EGFP images of the cortex injected with AAV7 expressing EGFP by the CBh, CMV, or CAG promoter. Scale bar, 100 μm. (**B**-**E**) Graphs showing ratios of respective EGFP-immunolabeled cell types to total EGFP-expressing cells by AAV7 vectors with the CBh, CMV, and CAG promoters. The dotted lines in the graphs are ratios of respective cell types to total cells present in the marmoset cortex (Figure S3). Error bars indicate S.E.M., and dots in the graph indicate the respective values for each of the individual marmosets. P values obtained using 1-way ANOVA with Tukey’s post hoc test were described in the graphs. n.d., not detected.

No or only minimal transduction of astrocytes in the marmoset cortex may be due to the small tropism of AAV vectors to marmoset astrocytes. However, we have previously shown efficient transduction of astrocytes in the marmoset cortex using AAV9 vectors carrying the astrocyte-specific cjGFAP promoter ^13^, suggesting that at least AAV9 capsid had tropism for marmoset astrocytes. To validate the tropism of AAV vectors for marmoset astrocytes, we injected the nine serotype vectors expressing EGFP by the astrocyte-specific human GFAP [hGFA(ABC1D)] promoter ^14^ (Figure 5A) into the marmoset cortex. Four weeks after the viral injection, the animals were sacrificed for fIHC. Confocal microscopy showed numerous astrocyte-like EGFP-expressing cells in all cortices injected with nine serotypes (Figure 5B). Subsequent fIHC analysis confirmed that most EGFP-labeled cells in cortical sections injected with any of the nine serotypes were immunolabeled for GFAP, confirming astrocyte transduction (Figure 5C, D). Although the difference was not statistically significant, AAV2 and AAV7 transduced astrocytes consistently with higher specificity (Figure 5D).

**Figure 5.**
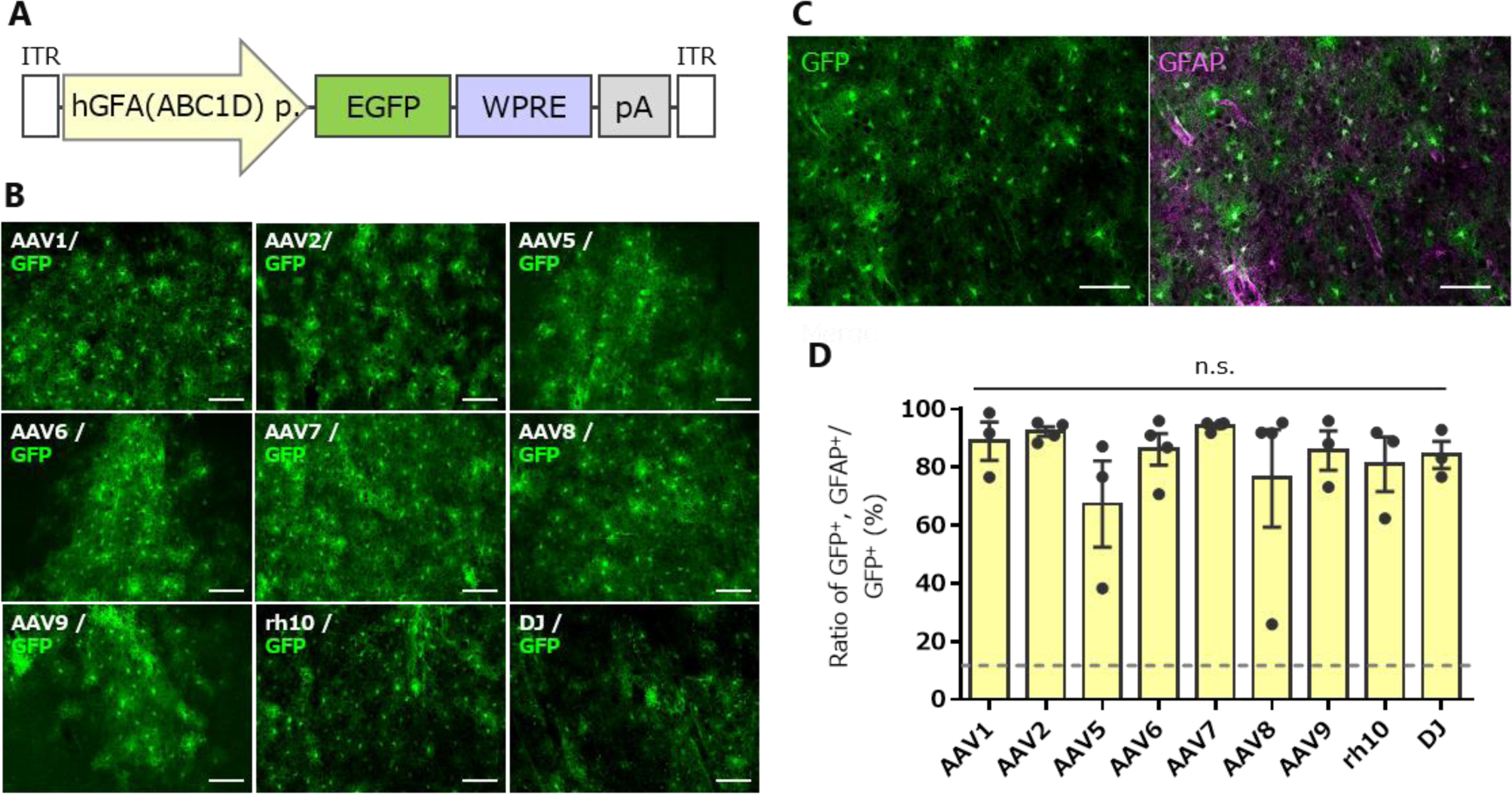
Efficient transduction of astrocytes by all nine serotype AAVs with the astrocyte-specific hGFA(ABC1D) promoter. (**A**) Schema depicting the AAV genome structure. (**B**) Immunofluorescent EGFP images of the cerebral cortex received injections of respective AAV vectors. Scale bar, 100 μm. (**C**) Representative image immunostained for EGFP alone (left) and merged image for EGFP and GFAP (right) after injection of the AAV2 vector. Scale bar, 100 μm. (**D**) Graph showing the percentage of GFAP-positive astrocytes to total EGFP-expressing cells 4 weeks after injection of the AAV vector as indicated. The dotted line in the graph shows a ratio of oligodendrocytes to total cells present in the marmoset cortex (Figure S3). Error bars indicate S.E.M., and dots in the graph indicate the respective values for each of the individual marmosets. n.s., not statistically significant by one-way ANOVA with Tukey’s post hoc test.

AAV5 and AAV8 transduced astrocytes variably with lower specificity (Figure 5D). We examined the results of individual marmosets and found that one marmoset (ID: H271, see Table 1) showed markedly low specificity for cortical astrocyte transduction when injected with AAV5 or AAV8. To identify cell types other than astrocytes that express EGFP, the cortical sections were immunostained with anti-Olig2 and anti-Iba1 antibodies. The results revealed that all EGFP-expressing non-astrocytes were immunolabeled for Olig2 (Figure S3), indicating that they were oligodendrocytes.

Since AAV5 carrying the CBh promoter also showed consistently higher transduction specificity for oligodendrocytes than many other serotypes (Figure 3C), we suppose that the AAV5 capsid may have tropism for oligodendrocytes. If AAV5 capsid is oligodendrocyte-tropic, AAV5 with the oligodendrocyte-specific promoter may achieve efficient and specific transduction of oligodendrocytes in the marmoset cortex. To validate this, we produced an AAV5 capsid vector expressing EGFP by the oligodendrocyte-specific mouse-derived MBP (mMBP) promoter (Figure 6A). AAVrh10 capsid was used as a control because this capsid-coated, CBh promoter-driven AAV is highly neuron-tropic and transduces oligodendrocytes less effectively compared with AAV5 (Figure 3B, C).

**Figure 6.**
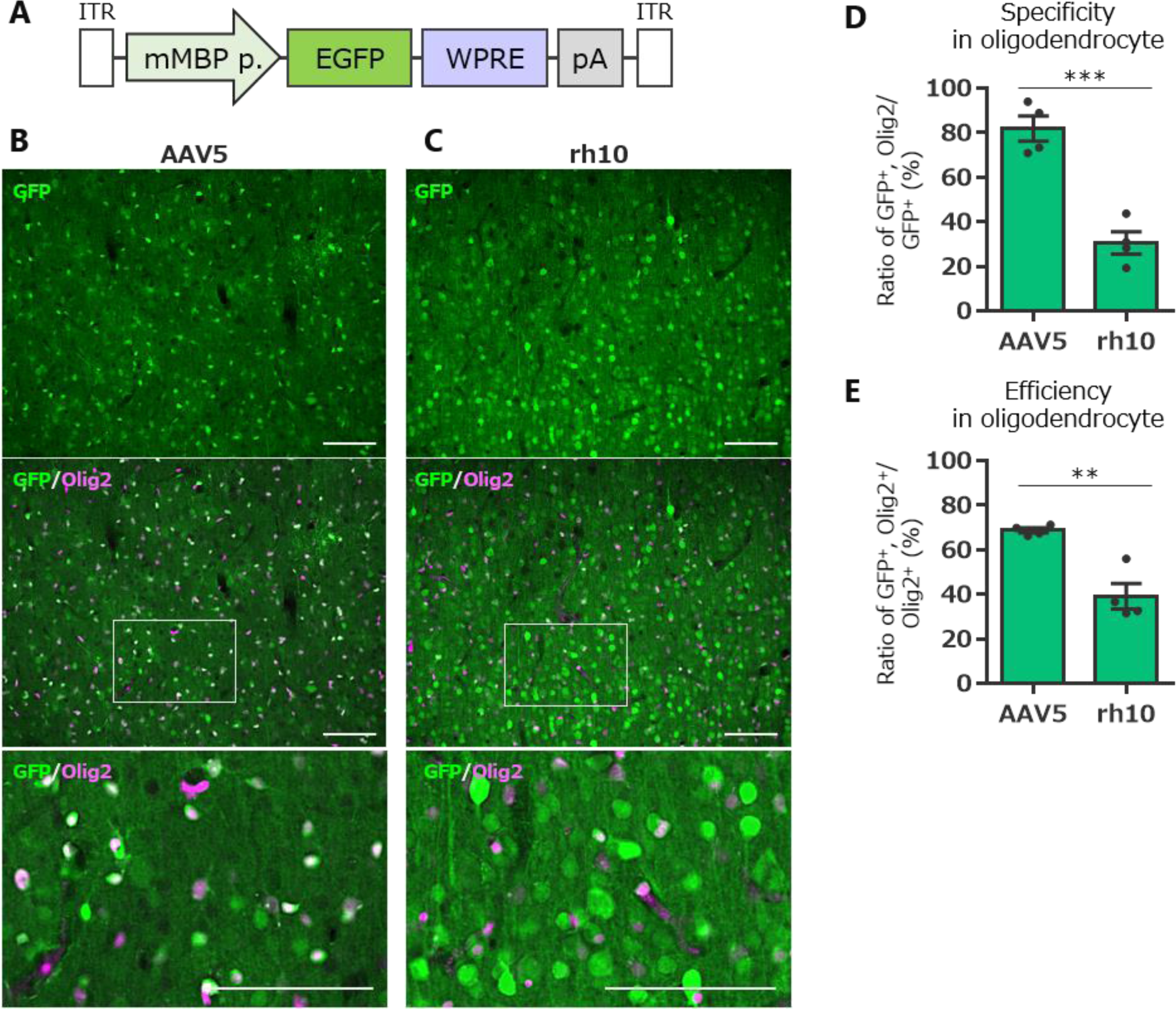
Selective and efficient transduction of oligodendrocytes in the marmoset cerebral cortex by AAV5 vectors with mouse myeline basic protein (mMBP) promoter. (**A**) Schema depicting the AAV genome structure. (**B**-**C**) Immunolabeled fluorescent images of EGFP in the cerebral cortex that received injection of AAV5 (**B**) or AAVrh10 (**C**) vectors expressing EGFP by the mMBP promoter. The middle immunofluorescence images present an overlay of immunolabeling for EGFP and the oligodendrocyte marker Olig2. The bottom images are magnifications of the boxed areas in the center images. Scale bar, 100 μm. (**D**-**E**) Summary graphs showing the specificity (**D**) and efficiency (**E**) of oligodendrocyte transduction. The dotted line in the graph indicates a ratio of oligodendrocytes to total cells present in the marmoset cortex (Figure S3). Error bars indicate S.E.M., and dots in the graph indicate the respective values for each of the individual marmosets. Asterisks indicate statistically significant differences between the AAV5 and AAVrh10. \*\**p* < 0.01, \*\*\**p* < 0.001 by student’s t-test.

Four weeks after the viral injection, animals were sacrificed for fIHC. Immunostaining of the cortical sections showed numerous EGFP-expressing cells co-immunolabeled for Olig2 in the marmoset that received injection of AAV5 (Figure 6B), in contrast to the much lower frequency of simultaneous immunolabeling of EGFP and Olig2 in the marmoset injected with AAVrh10 (Figure 6C). Quantitative results showed that over 80% of EGFP-expressing cells were Oig2-positive oligodendrocytes in marmosets injected with AAV5 (81.8 ± 5.7%, n = 4 marmosets), which were significantly higher than marmosets injected with AAVrh10 (30.6 ± 5.0%, n = 4 marmosets, \*\*\**p* = 0.0005 by student’s *t*-test) (Figure 6D). In addition, the transduction efficiency of oligodendrocytes was significantly higher in marmosets injected with AAV5 (68.8 ± 1.1%, n = 4 marmoset) than in marmosets injected with AAVrh10 (39.2 ± 5.7%, n = 4 marmoset, \*\**p* = 0.0023 by student’s t-test) (Figure 6E). Notably, although the oligodendrocyte-specific mMBP promoter was used, the specificity of oligodendrocyte transduction by AAVrh10 injection (∼31%) was less than the percentage of oligodendrocytes present in the marmoset cortex (∼42%, Figure S2).

## Discussion

In this study, we injected nine AAV serotypes expressing EGFP into the marmoset cerebral cortex and investigated the tropism of different cortical cell types in the marmoset brain. Although it did not reach a statistically significant level, AAV1, AAV9, AAVrh10, and AAV-DJ tended to cause widespread expression of EGFP. Subsequent immunohistochemistry showed that all serotypes with the CBh promoter expressed EGFP primarily in neurons. Among them, considering the spread of EGFP expression region, AAV9, AAVrh10, and AAV-DJ are suitable to efficiently express transgenes in marmoset cortical neurons. AAV7 and AAV2 stably expressed EGFP in astrocytes when combined with the astrocyte-specific hGFA(ABC1D) promoter, while AAV5 carrying the oligodendrocyte-specific mMBP promoter selectively expressed EGFP in oligodendrocytes.

The CBh, CMV, and CAG promoters are known to be constitutive promoters that are active in a variety of cell types ^15–17^. However, regardless of the serotypes used, these promoter-driven AAV vectors largely transduced neurons, with only a few transductions of glial cells in the cerebral cortex of marmosets. Notably, the CBh and CMV promoters did not transduce any astrocytes (Figs. 3D and 4C). Similar results were reported previously, which showed highly neurotropic transduction and almost no astrocyte transduction by direct injection of AAV8 or AAV9 with the CAG promoter in the marmoset brain ^18,19^. Thus, these constitutive CBh, CMV, and CAG promoters, which were delivered by parenchymal injection of AAV, worked specifically on neurons in the marmoset brain.

The results that AAV vectors with the so-called ubiquitously active CBh, CMV, or CAG promoter transduced no or only a few glial cells in the marmoset cortex do not indicate that the AAV serotypes tested had no ability to infect glial cells, because all nine serotypes efficiently transduced astrocytes when the astrocyte-specific hGFA(ABC1D) promoter was used, and AAV5 and AAVrh10 with the oligodendrocyte-specific promoter transduced oligodendrocytes (Figs. 5 and 6). Therefore, all nine AAV serotype capsids are tropic not only to neurons but also to astrocytes and oligodendrocytes. However, for microglia in marmoset cortex, it remains unclear whether the AAV serotypes used are unable to infect microglia or whether the promoter used is not activated in microglia.

Our previous experiments showed that intravenous injection of AAV9 expressing EGFP under the control of the CBh promoter caused efficient EGFP expression in astrocytes of the marmoset cerebral cortex ^20^. This contradicts the current result that the CBh promoter is not functional in marmoset cortical astrocytes. In this study, we injected AAV vectors directly into the marmosets’ cortex. This causes local tissue damage and cell death, leading to astrocyte activation. Thus, the CBh, CMV, and CAG promoter activities may be suppressed in reactive astrocytes.

Despite using the same oligodendrocyte-specific mMBP promoter, oligodendrocyte transduction specificity and efficiency differ widely and significantly between AAV5 and AAVrh10. Namely, AAV5 transduced oligodendrocytes with a high specificity of over 80%, whereas the specificity for oligodendrocytes by AAVrh10 was approximately 31%, which is less than the proportion of oligodendrocytes (∼42%) in total cortical cells (Figure 6E and Figure S2). This suggests that the tropism for oligodendrocytes differs greatly depending on the serotypes. Our results suggest that AAV5 is suitable for targeting marmoset oligodendrocytes.

Although the large differences in the specificity between serotypes as seen in oligodendrocytes were not observed in astrocytes, AAV2 and AAV7 stably transduced astrocytes with high specificity, and thus, AAV2 and AAV7 are thought to be suitable for targeting marmoset astrocytes. Like AAVrh10 with the oligodendrocyte-specific mMBP promoter, AAV5 with the astrocyte-specific hGFA(ABC1D) promoter transduced numerous non-astrocytes (Figure S3). These results suggest that the mMBP promoter and hGFA(ABC1D) promoter function as their respective target cell-specific promoters also in marmosets, but the cell type specificity in the marmoset brain is less than that in the mouse brain.

Here, using marmosets, we showed that nine AAV serotypes are capable of infecting neurons, astrocytes, and oligodendrocytes with distinct tropisms. It seems likely that the activities of the CBh, CMV, and CAG promoters are suppressed significantly in glial cells and function as neuron-specific promoters when AAV vectors are directly injected into the cerebral cortex. Astrocytes can be transduced with high specificity by AAV2 or AAV7 with the hGFA(ABC1D) promoter. Efficient oligodendrocyte transduction is achieved by AAV5 with the mMBP promoter. Therefore, selecting the appropriate promoter and optimal capsid is important to achieve target cell-specific transduction in the marmoset brain.

## Materials and Methods

### Animals

This study included 19 common marmosets (Calithrix jacchus) (summarized in Table 1). All marmosets are homebred at the Gunma University Bioresource Center. The animals were maintained in breeding rooms under controlled temperature (27–30 °C), humidity (25–45%), and light cycle (12 h each of light and dark) conditions. They could drink filtrated water, which was provided ad libitum. We fed 45–50 g of soaked monkey chow (CMS-1; CLEA Japan, Tokyo, Japan) with fruits, vegetables, or boiled chicken around noon, and marmoset-dumplings made by mixing CMS-1 soaked in hot water, honey, oligosaccharide, milk powder, vitamin supplement, lactobacillus powder, and gum arabic powder around three o’clock on a weekday afternoon. Cages and living space were suitable for GUIDE FOR THE CARE AND USE OF LABORATORY ANIMALS 8th edition. All efforts were made to minimize suffering and reduce the number of animals used, and all procedures regarding animal care and treatment were performed in accordance with guidelines approved by The Japan Neuroscience Society (‘Guidelines for experiments on primates in the field of neuroscience’) and the Institutional Committee of Gunma University (approval No. 20-053, 21-063, and 23-057).

### Construction of plasmids

The expression plasmids pAAV-CBh-EGFP-WPRE-HBGpA, pAAV-CMV-EGFP-WPRE-HBGpA, or pAAV-CAG-EGFP-WPRE-SV40pA were used as expression plasmids for constitutive expression of EGFP by the CBh promoter, CMV IE promoter, and CAG promoter, respectively^21–23^. These promoters were inserted into the expression plasmid pAAV just upstream of an EGFP gene at the restriction enzyme sites for XhoI and AgeI. The astrocyte-specific human-derived GFA(ABC1D) promoter from the pZac2.1-gfaABC1D-cyto-GCaMP6f gifted from Baljit Khakh (Addgene plasmid # 52925; http://n2t.net/addgene:52925; RRID:Addgene_52925) was amplified by KOD One PCR Master Mix (KMM-201; Toyobo, Osaka, Japan) using the following primers: 5’-ATGCTCTAGACTCGAGAACATATCCTGGTG-3’ and 5’-CATGGTGGCGACCGGTGCGAGCAGC-3’ to create the pAAV-hGFAP(ABC1D)-EGFP-WPRE-SV40pA^24^. The oligodendrocyte-specific mouse-derived MBP promoter was amplified from the pAAV-MBP-2xNLS-tdTomato gift from Viviana Gradinaru (Addgene plasmid # 104054; http://n2t.net/addgene:104054; RRID:Addgene_104054) by KOD One PCR Master Mix using following primers 5’-ATGCTCTAGACTCGAGTCCTTCCTGCTTAGGCCGTG-3’ and 5’-CATGGTGGCGACCGGTCCTCCGGAAGCTGCTGTGGG-3’ to create the pAAV-mMBP-EGFP-WPRE-SV40pA^25^. The PCR-amplified promoter fragments were inserted into the XhoI-AgeI site of the pAAV using Ligation high Ver.2 (LGK-201; Toyobo: hGFAP(ABC1D) or In-Fusion HD Cloning Kit (Takara Bio, Shiga, Japan: mMBP promoter).

The rep/cap plasmids pRC1 (TaKaRa Bio), pAAV2/5 (gifted from Melina Fan [Addgene plasmid # 104964; http://n2t.net/addgene:104964; RRID:Addgene_104964]), pAAV2/7 (gifted from James M. Wilson [Addgene plasmid # 112863; http://n2t.net/addgene:112863; RRID:Addgene_112863]), pAAV2/8 (gifted from James M. Wilson [Addgene plasmid # 112864; http://n2t.net/addgene:112864; RRID:Addgene_112864]) and pAAV2/AAVrh10 (gifted from James M. Wilson [Addgene plasmid # 112866; http://n2t.net/addgene:112866; RRID:Addgene_112866]) were obtained from Addgene. pAAV-DJ was purchased from Cosmo Bio (VPK-420-DJ; Cell Biolabs, San Diego, CA). pRC2-mi342 used to produce the AAV2/2 vector was a plasmid included in the AAVpro Helper Free System (Takara Bio). The AAV2/9 plasmid was kindly provided by James M. Wilson. To make rep/cap plasmid pAAV2/6, we replace the cap8 gene in pAAV2/8 (Addgene plasmid # 112864) with the cap6 gene in pRepCap6 (gifted from David Russell [Addgene plasmid # 110770; http://n2t.net/addgene:110770; RRID:Addgene_110770]). These gene engineering experiments were approved by the Institutional Committee of Gunma University (approval No. 20-018 and 23-056).

### Production of AAV vectors

Eight serotypes of AAV/CBh-EGFP vectors without AAV-2/CBh-EGFP and the AAV vectors with cell type-specific promoters were collected from the supernatant released outside the culture cells. Recombinant single-strand AAV vectors were produced using the ultracentrifugation method described in a previous paper were produced using HEK293-T cells (HCL4517; Thermo Fisher Scientific; Waltham, MA), as described previously^26^. Briefly, HEK293-T cells, which were cultured in Dulbecco’s Modified Eagle Medium (D-MEM; D5796-500ML, Merck, Darmstadt, Germany) supplemented with 8% fetal bovine serum (26140-079, Sigma-Aldrich) at 37 °C in 5% CO2, were transfected with three plasmids: an expression plasmid pAAV, pHelper (Stratagene, La Jolla, CA), and a rep/capsid plasmid using Polyethylenimine “Max” (24765-1; Polysciences, Inc., Warrington, PA, USA). Viral particles were harvested from the culture medium 6 days after transfection and were concentrated by precipitation with 8% polyethylene glycol 8000 (Merck) and 500 mM sodium chloride. The precipitated AAV vectors were re-suspended in D-PBS(-) and purified with iodixanol (OptiPrep; Serumwerk Bernburg AG, Bernburg, Germany) through continuous gradient centrifugation. The viral solution was further concentrated in D-PBS(-) using Vivaspin 20 (100,000 MWCO PES, Sartorius, Gottingen, Germany).

In addition, Dr. Hioki of the Brain/Minds viral vector core provided us with the AAV2/CBh-EGFP vector extracted from inside and outside the cells. AAV vector particles were produced and purified as previously described (10.1371/journal.pone.0169611, 10.1007/978-1-0716-1522-5_22). Briefly, pAAV-CBh-EGFP-WPRE-HBGpA and two helper plasmids, pBSIISK-R2C1 (10.1371/journal.pone.0169611) and pHelper (Merck; GenBank accession No: AF369965.1), were co-transfected into HEK293T cells (RCB2202; Riken BRC) using polyethylenimine (23966; Polysciences). Virus particles were purified from the cell lysate and supernatant using ultracentrifugation with OptiPrep (Serumwerk Bernburg AG) and concentrated using ultrafiltration with Amicon Ultra-15 (UFC903024; Merck).

The genomic titers of the viral vectors were determined by real-time quantitative PCR using Thermal Cycler Dice Real Time System II TP900 or III TP970 (Takara Bio Inc.) and Power SYBR Green PCR Master Mix (Thermo Fisher Scientific), using the primers 5’-CTGTTGGGCACTGACAATTC-3’ and 5’-GAAGGGACGTAGCAGAAGGA-3’, which targeted the WPRE sequence. The expression plasmid was used as a standard to plot for absolute quantitation. The produced AAVs were stored at 4°C for a few months or less and at −80°C for longer storage. AAV2 vectors were frozen and stored at −80°C.

### Cerebral cortical parenchymal viral administration

For immobilization of marmosets during parenchymal administration, we anesthetized marmosets with a cocktail of ketamine hydrochloride (20–25 mg/kg) and xylazine hydrochloride (4–5 mg/kg) and maintained the anesthetic state with isoflurane (2–2.5% in 60–70% O2, 1 L/min) using anesthesia apparatus (NARCOBIT-E(II), KN-1071; Natsume Seisakusho, Tokyo, Japan). SpO2 concentration and heart rate were monitored with a pulse oximeter (OLV-2700; Nihon Kohden Co., Tokyo, Japan). The marmoset was held in a brain stereotaxic instrument (SR-5C-HT; Narishige, Tokyo, Japan), and a thermal seat was used to maintain the body temperature of the marmoset. After the scalpel incision was made, a hole was drilled into the skull against the viral administration point using an electric drill (DC Power Pack C2012 and handpiece Minimo SD-101 attaching carbide cutter BC1403 or steel drill KA1001; Minitor Co. Tokyo, Japan). A 30-or 32-gage needle with a 1-2mm angled tip was used to confirm that the skull had been punctured and to simultaneously injure the meninges. The AAV solutions were loaded into a 33G Hamilton syringe (701SN 33G 2’’/PT3, 80308; Hamilton Co., Reno, NV), set into a microinjector (IMS-30; Narishige) attached to a stereotaxic instrument. The needle was inserted 1 mm below the base of the skull, and the AAV solutions were administered at a flow rate of 0.1 μL/min. After all viral injections were completed, the holes were plugged with medical-grade Aron Alpha A (Daiichi Sankyo Co., Tokyo, Japan) and sutured with a synthetic absorbable suture. Finally, to prevent the marmosets from scratching the sutures and incisions with their own fingernails, liquid adhesive plaster, Coloskin (Tokyo Koshi, Tokyo, Japan), was used to cover the sutures and wounds. The antibiotic ampicillin (5 mg) was administered for 5 days to prevent infection.

### Necropsy

Sacrifice was performed 4 weeks after the AAV injection. Marmosets were anesthetized with a cocktail of ketamine hydrochloride and xylazine hydrochloride for induction of anesthesia and isoflurane. Marmosets were perfused with 300 ml of cold 1 x PBS(-) containing 20 mM EDTA (311-90075, Nippon Gene, Tokyo, Japan) and fixed with 250 ml of cold 4% paraformaldehyde (PFA) in 1 x phosphate buffer (PB), and the brains were removed. GFP fluorescence on the brain surface was captured by fluorescence microscopy (VB-7010; Keyence, Osaka, Japan), and brains were postfixed in 4% PFA overnight.

### Fluorescent immunohistochemical analysis

Brain slices were prepared for GFP expression analysis. Trimmed except for the cerebrum and embedded in 2% agarose gel to make 100 µm thick sagittal sections using a microtome (VT1200S; Leica Microsystems GmbH, Wetzlar, Germany). Sections were stored at 4 °C in 1 x PBS(-) with NaN3 until use. Fluorescent immunohistochemistry (fIHC) in free-floating was performed to identify GFP expression in tissues and various kinds of brain cells. Tissues were quadruple fluorescently stained, including nuclear staining using NucBlue (Hoechst 33342). Tissue sections were reacted overnight at room temperature by immersion in following primary antibodies in blocking solution (2% Donkey Serum (S30-100ML, Merck), BSA (01862-87, Nacalai Tesque, Kyoto, Japan), 0.5% Triton X-100, 0.03% NaN3 in 1 x PB): rat monoclonal anti-GFP antibody (1:1,000; 04404-84; Nacalai Tesque, Kyoto, Japan), mouse monoclonal anti-NeuN antibody (1:1,000; MAB377; Merck), rabbit polyclonal anti-GFAP antibody (1:200; GFAP-Rb-Af800; Nittobo Medical, Tokyo, Japan), rabbit polyclonal anti-S-100β antibody (1:200; S100b-Rb-Af1000, Nittobo Medical), mouse monoclonal anti-Olig2 antibody (1:500; MABN50; Merck) and rabbit polyclonal anti-Iba1 antibody (1:500; 019-19741; Fujifilm Wako Chemicals, Tokyo, Japan). To visualize the bound primary antibodies, the sections were incubated for 3–4 hours at room temperature in the blocking solution containing the following secondary antibodies: Donkey anti-rat IgG Alexa Fluor Plus 488, Donkey anti-mouse IgG Alexa Fluor Plus 555, Donkey anti-rabbit IgG Alexa Fluor Plus 555, Donkey anti-rat IgG Alexa Fluor Plus 647 (1:2,000, Thermo Fisher Scientific). After the secondary antibody reaction, they were sealed in glass slides using ProLong Glass Antifade Mountant with NucBlue Stain (Thermo Fisher Scientific), cured, and stored at 4 °C. The primary and secondary antibody information used in this study is listed in Table 1.

Immunostained slices were photographed with a fluorescence microscope BZ-X800 (Keyence). The images used for cell counting were taken at each injection site using the same exposure time settings and sectioning function. All images for cell type counting were taken with a 20x objective and had an area of 0.394 mm2. On the other hand, a consolidated image of 0.672 mm2 was used for counting the number of cells of each endogenous cell type. The counting of cells on the images was done using the free software katikati counter (https://www.vector.co.jp/soft/win95/art/se347447.html).

### Statistics analysis

GraphPad Prism ver. 6 (GraphPad Software, San Diego, CA) was used for statistical analysis and output of graphic images. The analysis of variance among multiple groups was performed by a 1-way ANOVA with Tukey’s multiple comparison test. A student’s t-test was used to compare the results of the two groups. Each set of data was expressed as scatter plots with bar graphs. Bars indicated mean values, error bars indicated standard error of the mean (SEM), and black dots indicated data for each marmoset.

### Data availability

The datasets and programs generated for this study are available from the corresponding author upon request.

## Acknowledgments

The authors thank Asako Ohnishi, Nobue McCullough, and Chieko Miyazawa for AAV1, 5, 6, 7, 8, 9, rh10, DJ vector production, Prof. Hiroyuki Hioki at Juntendo University for AAV2 vector production, Motoko Uchiyama, Minako Noguchi, and Yoshiko Nomura for raising the marmosets, and Junko Sugi for immunohistochemistry. This research was partially supported by the program for Brain Mapping by Integrated Neurotechnologies for Disease Studies (Brain/MINDS) from the Japan Agency for Medical Research and Development (AMED) (JP20dm0207057, JP21dm0207111, and JP21dm0207112), JSPS KAKENHI (grant numbers 23H02791, 19K06899, and 22K06454, respectively), and Gunma University for the promotion of scientific research.

## Author contributions

H. H. supervised the study. Y.M., A.K., and H.H. designed the experiments. Y.M., Y.F., and A.K. performed experiments. Y.M. prepared the original drafts. All the authors have read and approved the final version of the manuscript.

## Declaration of interests

The authors declare no competing interests.

## Notes

### Competing Interest Statement

The authors have declared no competing interest.

