## Supplementary figures for "Optimal AAV capsid/promoter combinations to target specific cell types in the common marmoset cerebral cortex"

### Figure S1

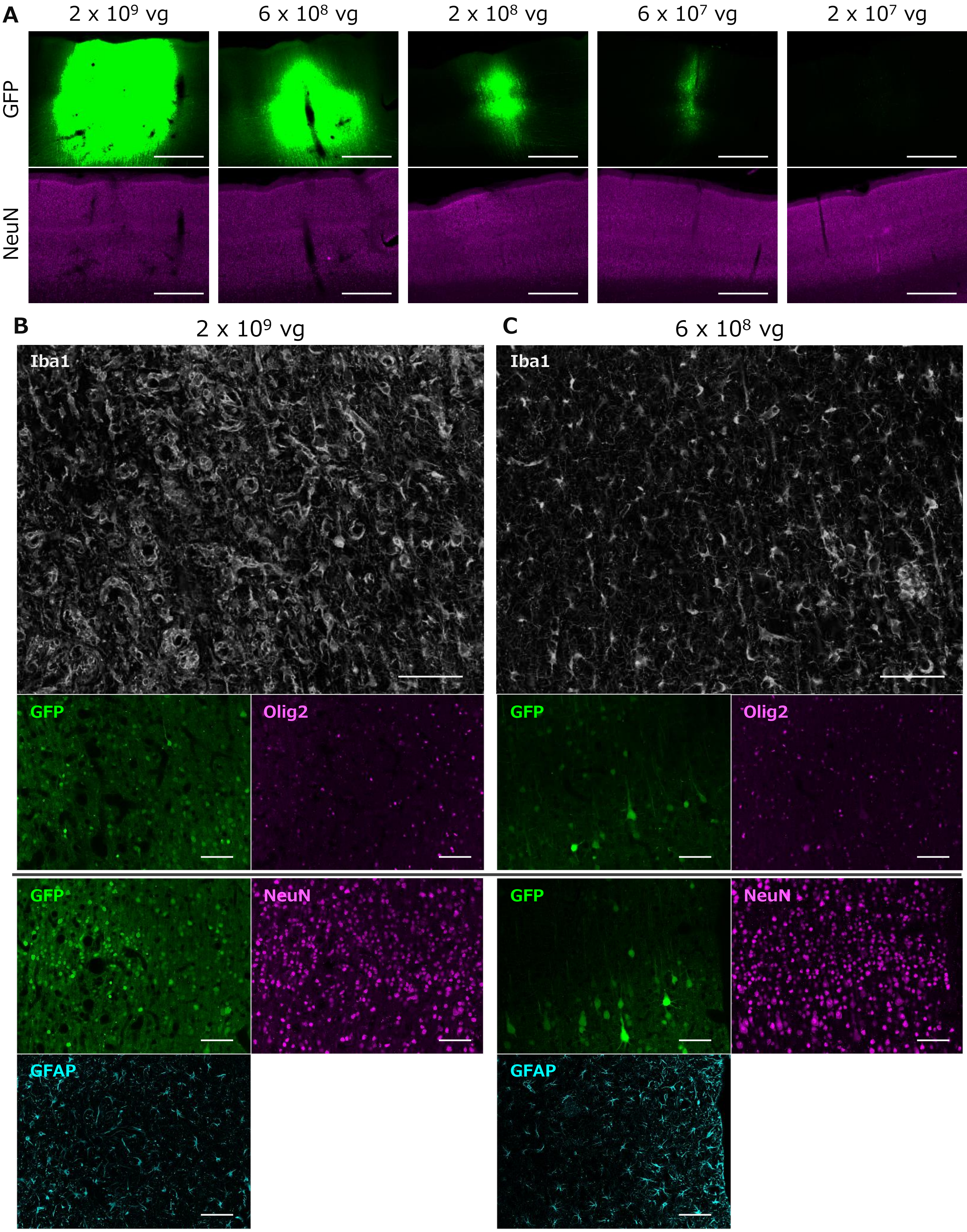
A
2 x 10^9 vg
6 x 10^8 vg
2 x 10^8 vg
6 x 10^7 vg
2 x 10^7 vg
GFP
NeuN
B
2 x 10^9 vg
C
6 x 10^8 vg
Iba1
GFP
Olig2
NeuN
GFAP

**Figure S1.** Figure S1. Spread of AAV vectors and microglial activation by direct injection of AAV vectors into the marmoset cerebral cortex

**(A)** 3-fold dilution series of AAV2 vectors expressing EGFP by the CBh promoter was injected into the marmoset cerebral cortex. Immunofluorescent images of EGFP (upper) and NeuN (lower) in the marmoset cortex 4 weeks after AAV injection at the doses indicated. Increasing the amount of virus injected expanded the area of EGFP expression. However, no obvious neuronal loss was observed even when the highest dose was injected. **(B-C)** Injection of AAV2 at $2.0 \times 10^9$ vg caused considerable activation of microglia **(B)**, while no apparent activation was observed by a lower amount of AAV2 at $6.0 \times 10^8$ vg or lower **(C)**. The top two large images show microglia immunolabeled for Iba1. The smaller images below are immunohistochemistry for GFP, Olig2, NeuN, and GFAP, as labeled on the top left of each image. The images above and below the gray line in the center were obtained using different tissue sections. Scale bar, 100 μm.

### Figure S2

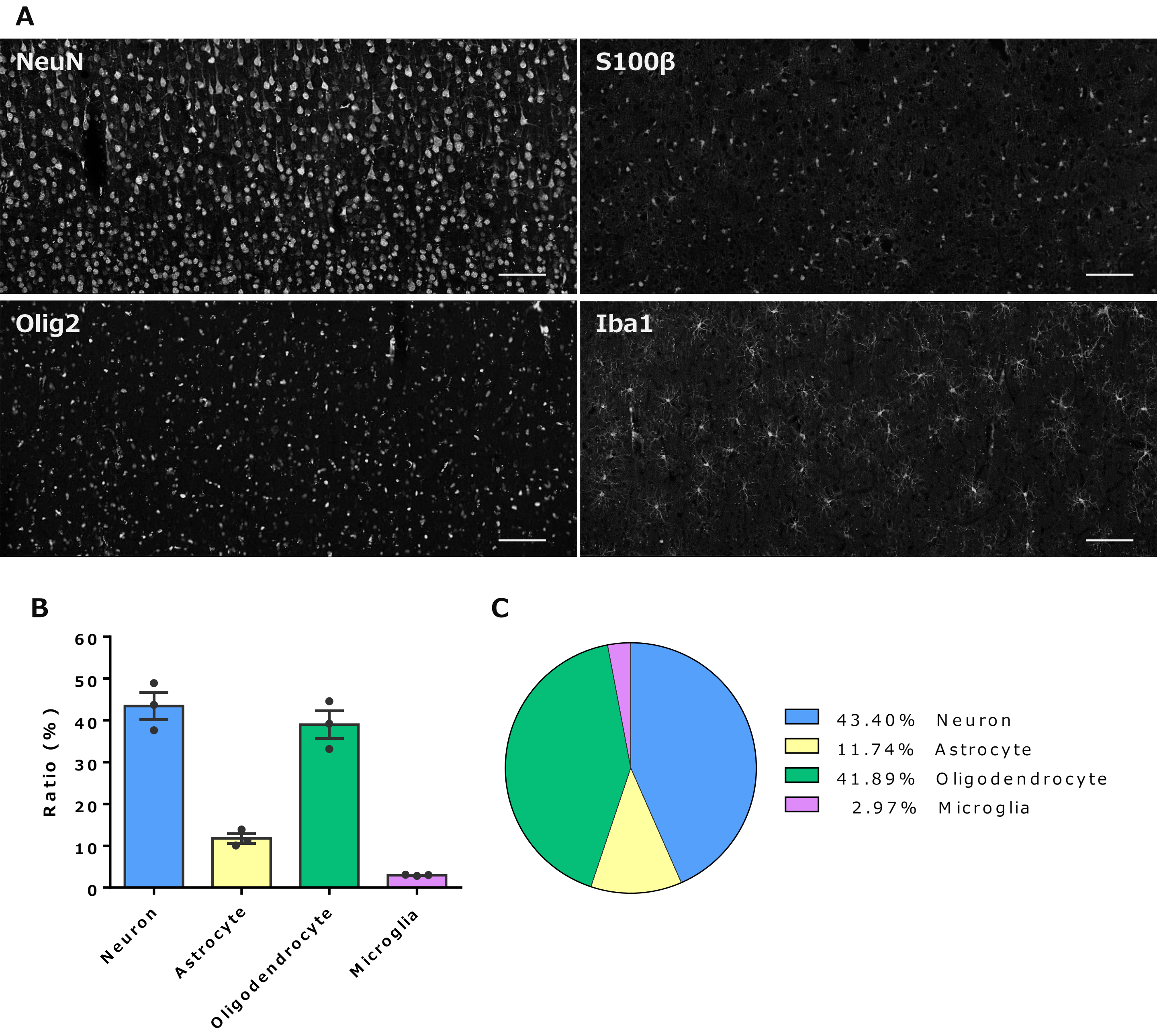


**Figure S2.** The proportions of different cell types endogenously present in the marmoset cerebral cortex were assessed immunohistochemically

(**A**) Representative fluorescent images immunolabeled for NeuN, S100β, Olig2, and Iba1. Scale bar, 100 μm. (**B**, **C**) Summary bar graph (**B**) and pie chart (**C**) showing the percentages of neurons, astrocytes, oligodendrocytes, and microglia. Error bars indicate S.E.M., and dots in the graph indicate the respective values for each of the individual marmosets.

#### Figure S3

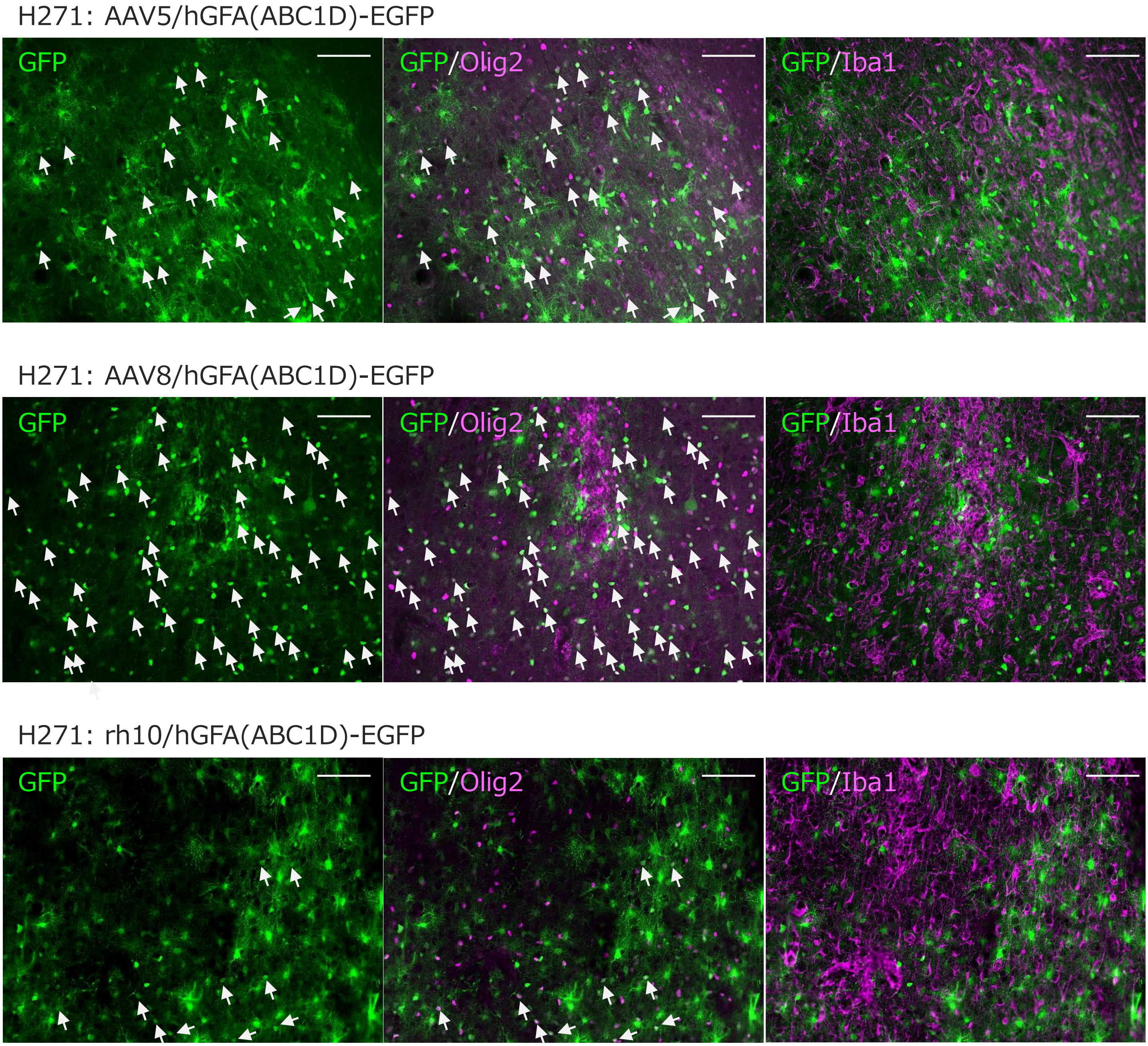


**Figure S3.** Examples of extensive oligodendrocyte transduction by the astrocyte-specific hGFA(ABC1D) promoter in the case of some AAV capsids

Immunofluorescent images of the cerebral cortex from marmoset (ID: H271, see Table 1) 4 weeks after injection of AAV5 (top), AAV8 (middle), and AAVrh10 (bottom) expressing EGFP by the astrocyte-specific hGFA(ABC1D) promoter. The cerebellar sections were triple immunostained for EGFP, Olig2, and Iba1. Note that numerous GFP-expressing cells were co-immunolabeled for Olig2 (arrows) when AAV5 and AAV8 were used, indicating extensive transduction of oligodendrocytes despite employing the astrocyte-specific promoter. Scale bar, 100 μm.
